# RNA regulates repeat-associated non-AUG (RAN) translation initiation in *C9orf72* FTD/ALS

**DOI:** 10.1101/2025.10.09.681477

**Authors:** Rosslyn Grosely, Shizuka Yamada-Hunter, Antonio V. Puglisi, Michael Z. Palo, Carlos Alvarado, Christopher P. Lapointe, Aaron D. Gitler, Joseph D. Puglisi

**Author notes:** Corresponding author: Joseph D. Puglisi.

## Abstract

Repeat-associated non-AUG (RAN) translation synthesizes protein in the absence of a cognate AUG start codon ^1,2^In frontotemporal dementia and amyotrophic lateral sclerosis, a GGGGCC (G_4_C_2_) repeat expansion in an intron of *C9orf72* leads to synthesis of neurotoxic dipeptide-repeat proteins, underscoring the need to understand the mechanism of *C9orf72* RAN translation^1-5^. RNA sequence and structure have been implicated, but how they direct *C9orf72* RAN translation, particularly the rate-limiting, multi-step initiation phase, remains unclear^6-10^. We applied single-molecule biophysics to a reconstituted human translation initiation system and tracked fluorescently labeled ribosomes and initiation factors in real time. We show that RNA G_4_C_2_ repeats and sequence context alter initiation factor dynamics after ribosomal scanning, generating a kinetic bottleneck in the commitment to initiate at a near-cognate CUG start codon. Our model of *C9orf72* RAN translation provides a mechanistic framework for how repeat expansions change underlying translation dynamics and may be broadly relevant to other disorders that involve RAN translation.

## Introduction

Amyotrophic lateral sclerosis (ALS) and frontotemporal dementia (FTD) are two fatal neurodegenerative diseases^11^. Mutations in the *C9orf72* gene are the most common genetic cause of ALS and FTD (C9FTD/ALS)^4,12^. The *C9orf72* gene contains a polymorphic hexanucleotide repeat, GGGGCC, located in an intron. The repeat tract length in unaffected individuals, although variable, is typically between five and ten repeats and almost always fewer than 23 repeats. In ALS and FTD, the hexanucleotide repeat tract is expanded to hundreds or even thousands of repeats. This repeat expansion may cause disease by several different mechanisms^13^. They include loss-of-function through haploinsufficiency^14^ and RNA toxicity through sense or antisense repeat-containing transcripts sequestering RNA-binding proteins^15^. Disease may also arise due to a non-conventional form of translation called repeat-associated non-AUG translation (RAN translation)^1-3^.

RAN translation of the RNA repeats yields dipeptide repeat proteins (DPRs) that are aggregation prone, accumulate in the nervous system, and are toxic to cells, contributing to FTD/ALS disease pathogenesis^16-18^. These DPRs include glycine-alanine (GA), glycine-proline (GP), glycine-arginine (GR), proline-arginine (PR), and proline-alanine (PA) and are translated from cytoplasmic repeat-containing RNAs that do not contain a canonical AUG start codon^1-3^. How this unusual translation initiation pathway in C9FTD/ALS occurs remains unclear, requiring analysis of the sequence features promoting RNA translation and the identification of regulators to illuminate molecular mechanisms^19^.

Eukaryotic translation initiation is a tightly regulated, multi-step process to assemble the 80S ribosome at the AUG start codon of an open reading frame (ORF)^20,21^. The eIF4F complex, composed of the cap-binding protein eIF4E, the RNA helicase eIF4A, and a scaffolding protein eIF4G, binds the 5^′^ m^7^G cap of the mRNA and loads 43S initiation complex, which consists of the 40S ribosomal subunit, eIF1, eIF1A, eIF3, and the eIF2-GTP-Met-tRNA_i_ ternary complex^22-24^. Following recruitment, the 43S complex scans single-stranded mRNA directionally 5^′^ to 3^′^ in an ATP-dependent manner, mediated by eIF4A and its cofactor eIF4B^25^. Scanning continues until the ribosome reaches an AUG codon^26-28^. Base-pairing between the anticodon loop of 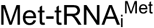 and the AUG start codon triggers a conformational rearrangement of the 40S subunit to a scanning-arrested state, initial dissociation of eIF1, and stabilization of 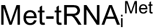 the P_in_ state^29-32^. As eIF1 then transiently rebinds the initiation complex, eIF5 competes to bind the complex^33-35^ and commit to the start site by stimulating eIF2 GTP hydrolysis^34,36-39^. These coupled steps expose the eIF5B-binding sites on Met-tRNAi and the ribosome allowing the GTPase eIF5B to bind and catalyze joining of the 60S ribosomal subunit^31,40-43^. Upon 60S subunit joining, eIF5B rapidly hydrolyzes GTP and dissociates leaving a fully assembled 80S initiation complex positioned at the AUG start codon, establishing the reading frame for translation elongation^44-46^.

In C9FTD/ALS, RNA structural elements present multiple obstacles to canonical translation initiation and subsequent synthesis of DPRs^47,48^. Unlike canonical mRNAs, *C9orf72* mRNAs with large intronic G_4_C_2_ repeats aberrantly retain the intron in the mature mRNA transcript^49-51^ and lack an upstream AUG start codon. These aberrant transcripts instead utilize utilizes a near-cognate CUG codon 24 nucleotides (nts) upstream of the first G of the repeats^52,53^. The G_4_C_2_ repeats have can form stable RNA secondary structures, particularly G-quadruplexes (GQs) or large hairpins that typically inhibit canonical translation^6,54-56^. However, in the context of C9FTD/ALS, these RNA elements paradoxically promote DPR synthesis^54,57-59^.

How G_4_C_2_ repeats promote translation is an open question. One hypothesis is that structured G_4_C_2_ repeats as an internal ribosome entry site (IRES), allowing for ribosome recruitment independent of the 5’ cap structure^16,53,58,60^. Alternatively, structured G_4_C_2_ repeats may act as a steric barrier to slow the scanning ribosome during 5’ cap-dependent initiation leading to a pause at the CUG codon; initiation at the near-cognate CUG would arise from an increased residency time of ribosomal initiation complex^61,62^. Consistent with these hypotheses, disrupting RNA structure reduces RAN translation efficiency, suggesting a functional role for structured RNA elements in promoting initiation^55,63^. Ribosome profiling data reveal frequent ribosomal stalling within repeats, further implicating the repetitive sequence and structure of G_4_C_2_ repeats as key contributors to aberrant translation^59,64^. Understanding how translation machinery engages these pathogenic RNAs, specifically how RNA structure facilitates RAN translation initiation is key to understanding C9FTD/ALS pathogenesis.

Despite its crucial role in C9FTD/ALS pathology, the mechanism of RAN translation initiation remains unknown. To address this, here we combined *in vitro* bulk and real time single-molecule assays to determine how RNA structural elements modulate the dynamics and efficiency of *C9orf72* RAN translation initiation.

## Results

To investigate how RNA structure regulates RAN translation in C9FTD/ALS, we used mRNA reporters containing tandem G_4_C_2_ repeats flanked by 120 and 105 nt of native *C9orf72* intronic 5^′^ and 3^′^ sequence, respectively (Fig 1a; Fig S1a -e). Downstream of the 3^′^ intronic sequence and in frame with the first G of the repeats, the reporter encodes a nanoLuc luciferase (nLuc) open reading frame that lacks an AUG start codon and ends in a poly(A) tail. Cell and animal models of C9 ALS RAN translation indicate 60 tandem repeats are sufficient to reproduce C9 ALS pathology^65-67^. We confirmed these mRNAs recapitulate *C9orf72* RAN translation by measuring nLuc activity after *in vitro* translation in HeLa cytoplasmic extract (Fig S1a-e). In-frame *C9orf72* RAN translation was ∼70% lower than a control mRNA and accounted for ∼90% of total *C9orf72* RAN translation across the three reading frames (Fig.S1b). As expected, mutation of the near-cognate CUG codon 24 nt upstream of the G_4_C_2_ repeats to a CCC inhibited translation (Fig.S1d). We thus focused all subsequent experiments on the in-frame RAN translation mediated by the CUG start site.

**Figure 1.**
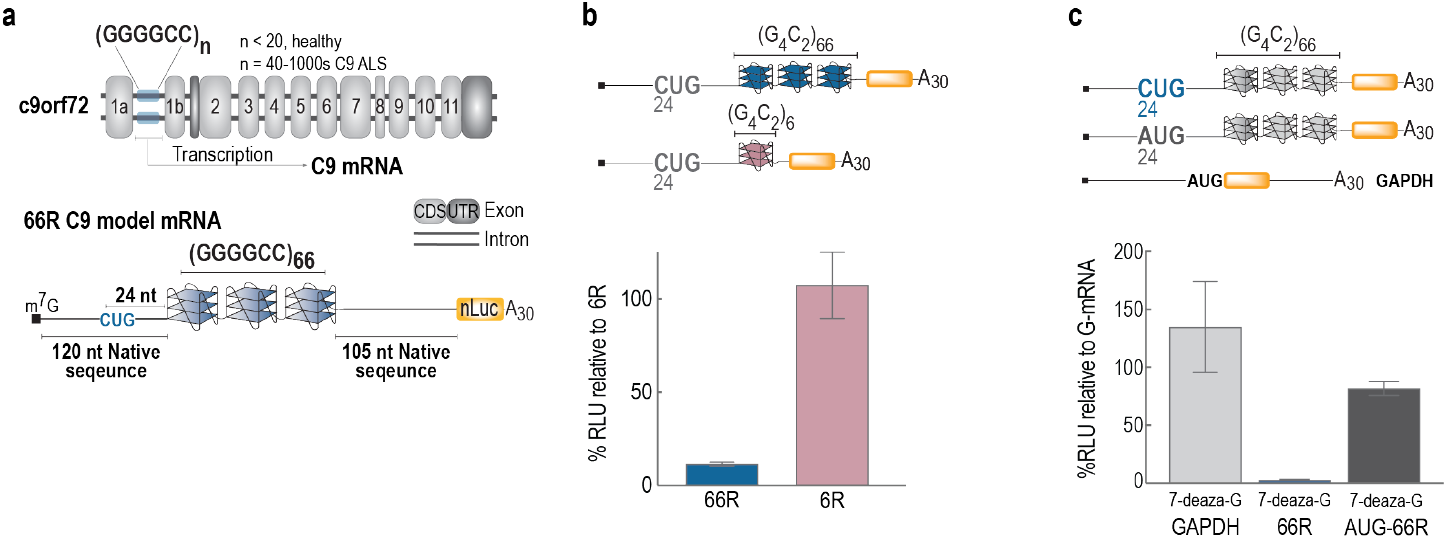
RNA structure modulates C9orf72 RAN translation in vitro. **a**.Schematic of the *C9orf72* gene locus illustrating the intronic G_4_C_2_ repeat expansion (blue) that is retained in cytoplasmic mRNA. The model reporter mRNA contains 66 G_4_C_2_ repeats (66R), a 5^′^ cap (m^7^G), a poly(A) tail (A_30_), and a nanoLuciferase (nLuc) open reading frame lacking an AUG start codon. **b-c**. Schematic of mRNAs and relative nLuc activity following in vitro translation of reporters in HeLa cytoplasmic extract. Data represent mean ± s.d. of biological replicates. **b**.RAN translation decreases with G_4_C_2_ repeat length. nLuc activity of the 66R reporter relative to the 6 G_4_C_2_ repeat (6R) reporter. (n = 3). **c**.RAN translation affected by G-quadruplex (GQ) formation. nLuc activity of 7-deaza-Guanine mRNA relative to mRNA Guanine. (n = 6). GAPDH serves as a negative control; AUG-66R harbors a CUG-to-AUG substitution.

### C9orf72 RAN translation initiation requires RNA structure

To define the role of G_4_C_2_ repeats in *C9orf72* RAN translation, we examined how repeat length and structure influence translation. Reducing G_4_C_2_ repeats from 66 to 2 eliminated *C9orf72* RAN translation, while 6 repeats stimulated RAN translation, yielding ∼9-fold higher nLuc activity than 66 repeats (Fig S1a; Fig 1c). Formation of an RNA G_4_C_2_ G-quadruplex (GQ) minimally requires 3 repeats and readily forms with 6 repeats, suggesting RAN translation of the 6-repeat mRNA is due to GQ structure^6,54^. To examine the role of GQ structure in *C9orf72* RAN translation, we in vitro transcribed mRNAs with 7-deazaGTP, a GTP analog that disrupts Hoogsteen base pairing required for GQ formation^68^. Translation of the 66 G_4_C_2_ repeat mRNA containing 7-deazaG reduced translation by 97 ± 0.5% while it enhanced translation of the control GAPDH by 35 ± 39% (Fig. 1b). By contrast, 7-deazaG had little effect when the CUG start codon upstream of the 66 G_4_C_2_ repeats was replaced with an AUG codon (reduced by 10 ± 6%). These findings suggest GQ structure promotes C9orf72 RAN translation, likely by facilitating initiation at the upstream CUG codon.

### C9orf72 RAN mRNA perturbs initiation kinetics and efficiency

To determine how pathogenic G_4_C_2_ repeats mediate *C9orf72* RAN translation, we monitored the ribosomal subunits and key eIFs directly as they progressed through the entire initiation pathway in real-time using our reconstituted human translation initiation system and single-molecule fluorescence microscopy. As a comparison to *C9orf72* RAN translation, we selected β-globin mRNA, which we previously examined and yields efficient initiation in our translation assays (Fig S1g;^35,46^). In all single-molecule experiments, 5^′^-m^7^G capped mRNAs were tethered to imaging surfaces and pre-incubated with eIF4F proteins and ATP to prepare the mRNAs for initiation, and then we started the measurement adding pre-formed 43S initiation complexes, additional initiation factors, the 60S subunit, and GTP. Fluorescently labeled molecules were tracked during initiation in real time. In this set of experiments, Cy3-43S (green) initiation complexes (Cy3-40S subunit, 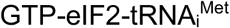, eIF1, eIF1A, eIF3, and eIF5), Cy3.5-eIF5B (yellow), and Cy5.5-60S (purple) were monitored (Fig 2a). Förster resonance energy transfer (FRET) between the 40S subunit Cy3 (donor) and 60S subunit Cy5.5 (acceptor; purple) fluorophores was used to identify 80S ribosome assembly. The association times (time between binding events) and lifetimes (duration of the binding event) were used to derive the reported mean times (see Table S1 for all rates and rate constants). In the kinetic plots, the mean time corresponds to the time at which the curve reaches y ≈ 0.63.

**Figure 2.**
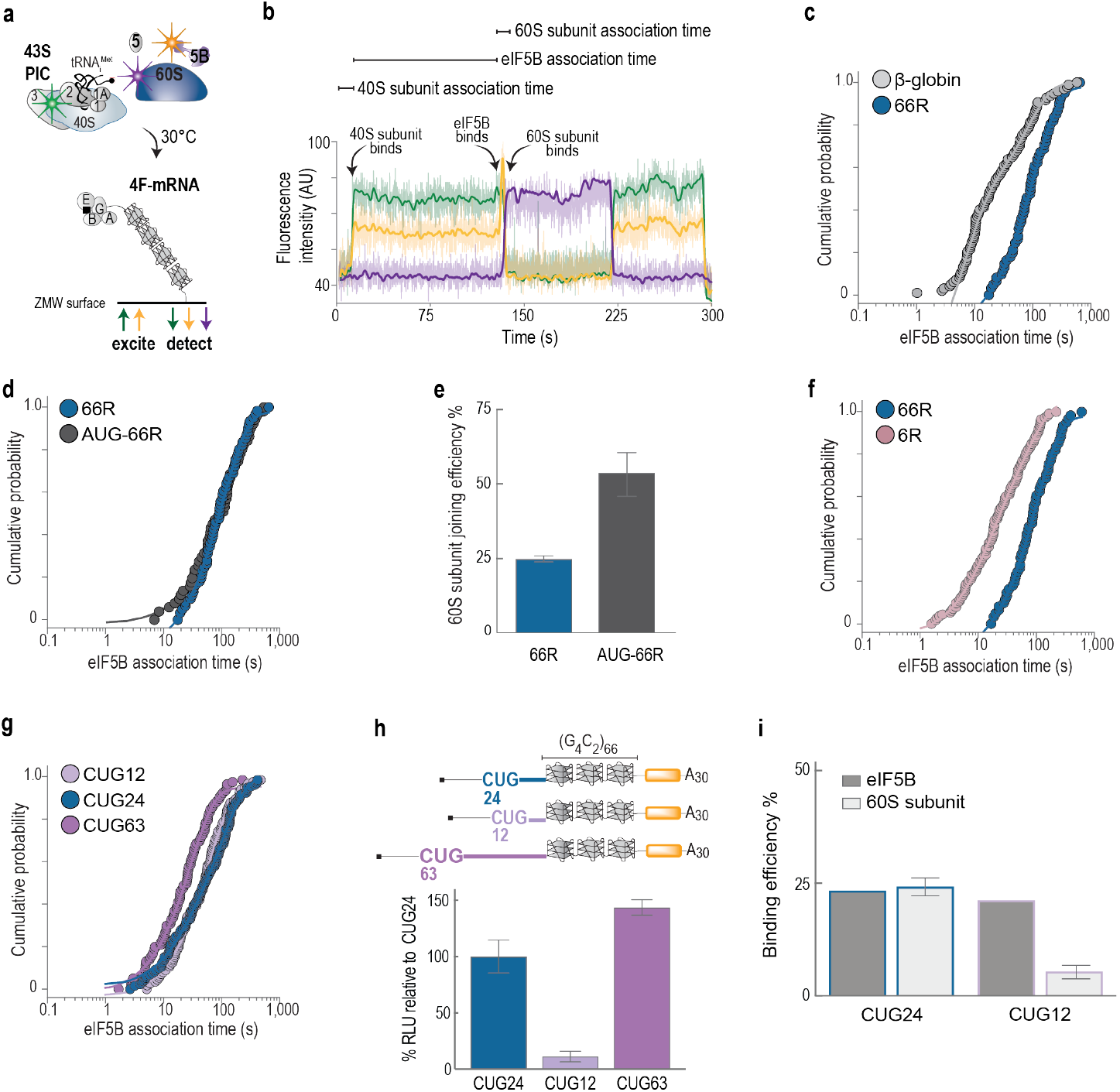
*C9orf72* RAN translation has altered initiation dynamics. **a**. Schematic of the real-time single-molecule human translation initiation assay. **b**. Representative single-molecule fluorescence trace tracking Cy3-40S subunit (green), Cy3.5-eIF5B (yellow), and Cy5.5-60S (purple). Ribosome assembly is marked by Cy3-Cy5 FRET. Molecular binding events are annotated in the example. **c–d**. eIF5B association time is delayed during *C9orf72* RAN translation initiation. The eIF5B association time for initiation on the 66 G_4_C_2_ repeat (66R) mRNA is compared to initiation on β-globin (**c**), the delay in eIF5B during *C9orf72* RAN translation initiation is independent of start codon identity. AUG-66R mRNAs carries a CUG-to-AUG mutation (**d**). Data are fit to an exponential function (lines) used to derive mean times and rates (Table S1). **e**. Start-codon identity modulates 60S subunit joining efficiency, defined as the fraction of eIF5B-bound molecules that proceed to 60S subunit joining. Data represent mean ± s.d. **f**. Decreasing the number G_4_C_2_ of repeats from 66 to 6 (6R) alleviates eIF5B perturbation. **g**. eIF5B association time is modulated by proximity of repeats to the CUG codon. Data are fit to exponential functions (lines on plots) used to derive mean times and rates (Table S1). **h**. *C9orf72* RAN translation level is modulated by the distance between the CUG codon and G_4_C_2_ repeats. nLuc activity was measured for reporters with 12 nt (CUG12), 24 nt (CUG24), or 63 nt (CUG63) spacing between CUG start codon and G_4_C_2_ repeats. Two nt substitutions were placed upstream of the CUG to insert stop codons in the +1 and +2 reading frames. (see Sup Table 2 for sequences). Data represent mean ± s.d of biological replicates (n = 4). **i**. Repeat proximity alters eIF5B and 60S joining efficiency. eIF5B efficiency is defined as the fraction of 40S-bound complexes that proceed to eIF5B association; 60S subunit joining efficiency is the fraction of eIF5B-bound complexes that proceed to 60S subunit joining. Data represent mean ± s.d.

The steps of RAN translation initiation followed the same order observed on β-globin mRNA, but with a striking delay. The 43S initiation complex loaded onto the *C9orf72* RAN mRNA with 66 G_4_C_2_ repeats and β-globin mRNA with similar kinetics, and both were dependent on eIF4F activity (Fig S2a-b) suggesting *C9orf72* initiation is 5’ cap-dependent initiation. In stark contrast, the delay between loading of the 43S initiation complex and eIF5B association lengthened by more than 5-fold on the RAN mRNA to 111 ± 3.5 s (Fig 2c). Once eIF5B bound, the timing of 60S subunit joining and subsequent time of eIF5B departure matched the timing on β-globin (Fig S2c) Thus, 66 G_4_C_2_ repeats mediate translation initiation, but our single-molecule findings indicate that the repeats dramatically inhibit and delay eIF5B binding, which ultimately limits successful formation of translation competent ribosomes on the mRNA.

### The repeats and their position regulate C9orf72 RAN translation

To determine whether the delay in eIF5B binding was due to the 66 G_4_C_2_ repeats or the native CUG start site, we replaced the CUG with a canonical AUG. We observed a near-identical delay until eIF5B bound (125 ± 9.0 s; Fig 2d), but joining of the 60S subunit was ∼2-fold more efficient (54 ± 10% compared to 25 ± 2%; Fig 2e). This finding suggests that the AUG site funneled more eIF5B-bound complexes to successful initiation, which we confirmed using our bulk translation assay (Fig S1d). In parallel, reduction of the number of G_4_C_2_ repeats from 66 to 6 downstream of the CUG allowed eIF5B to bind 3-fold more rapidly (32 s ± 1.7 s; Fig 2f), which agrees with its enhanced translation in bulk translation assays (Fig 1c). Thus, the CUG near-cognate start site and G_4_C_2_ repeats synergistically modulate *C9orf72* RAN translation but do so by targeting different steps of the initiation process.

If G_4_C_2_ repeats act as a barrier that positions the ribosome at the at the CUG start site, then spacing between CUG and G_4_C_2_ repeats should be critical. When positioned at the start codon, the leading side of the 40S ribosome covers 16-18 nt downstream, placing the G_4_C_2_ repeats ∼6-8 nt away from the ribosome^24,69^. To test the effects of G_4_C_2_ repeat proximity on *C9orf72* RAN translation initiation, we varied the distance between the CUG start codon and the 66 G_4_C_2_ repeats and monitored eIF5B binding kinetics. Increasing the distance between the CUG start site and the G_4_C_2_ repeats from 24 nt to 63 nt accelerated eIF5B association by nearly 3-fold (36 ± 1.2 s) (Fig 2g). Consistent with more rapid eIF5B binding kinetics, the increased distance enhanced RAN translation by 44 ±14% in our bulk assays (Fig 2h). By contrast, positioning the CUG start site 12 nt upstream of the repeats (closer) had limited effect on eIF5B binding kinetics (Fig 2g). However, the proximity of the CUG start codon to the repeats reduced the efficiency of 60S subunit joining to 7 ± 2% (Fig 2i). The inhibitory effect of 12-nt spacing suggests that the G_4_C_2_ repeat structures can sterically hinder 60S subunit. Consistently, reducing the spacing to 12 nt inhibited translation by 89 ± 5% in our bulk assay (Fig 2h). Our findings indicate that distance between the CUG start site and G_4_C_2_ repeats can modulate *C9orf72* RAN translation initiation.

The results presented above indicate that long, pathogenic-length G_4_C_2_ repeats natively positioned downstream of the CUG start site allow eIF4F-dependent initiation but substantially delay the process after the 43S initiation complex loads onto the mRNA. The G_4_C_2_ repeats likely function beyond facilitating CUG initiation as their length and position affect *C9orf72* RAN translation initiation dynamics. Steric hinderance of eIF5B association by the G_4_C_2_ repeats could explain these observations. Alternatively, the observed perturbations to eIF5B association could result from an upstream rate-limiting step, such as scanning or start codon recognition.

### Kinetics of scanning and start codon recognition are unaffected by G_4_C_2_ repeats

To determine if the G_4_C_2_ repeats modulate scanning, we monitored eIF1 directly during *C9orf72* RAN translation initiation. eIF1 stably binds the initiation complex during the scanning step, but transitions to a transient rebinding mode once the start site has been reached^35^. Therefore, in single-molecule assays, the time between 43S initiation complex loading on the mRNA and the first dissociation of eIF1 represents the time it takes for the scanning ribosome to reach the start codon. We assembled the 43S initiation complex as above but instead used eIF2-Cy3-tRNA_i_-GTP ternary complex (green), Cy5-eIF1 (red), and unlabeled 40S subunits. We monitored tRNA_i_-Cy3 (and thus the 43S complex) and eIF5B-Cy3.5 directly, and eIF1-Cy5 via tRNA_i_-to-eIF1 FRET during translation initiation (Fig 3b).

**Figure 3.**
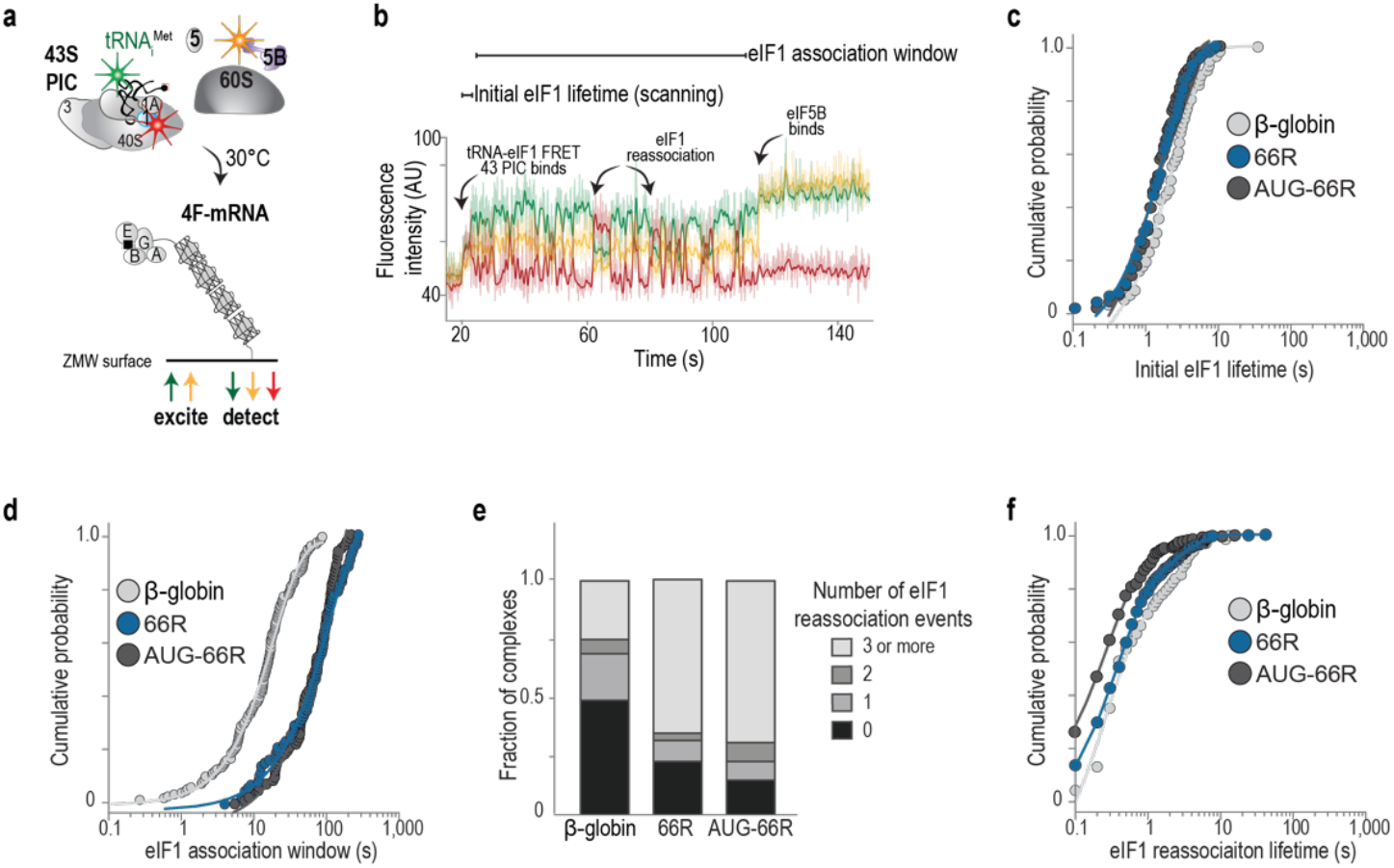
eIF1 dynamics are altered after scanning during C9 RAN translation initiation. **a**. Schematic of the single-molecule scanning and start codon recognition assay. **b**. Representative trace monitoring Cy3-tRNA_i_ (green), Cy5-eIF1 (red), and Cy3.5-eIF5B (yellow). eIF1association is tracked via Cy3-Cy5 FRET. Molecular biding events are notated. **c–d, f**. Cumulative probability plots showing eIF1 binding dynamics on the 66 G_4_C_2_ repeat (66R), the 66R mRNA with a CUG to AUG mutation (AUG-66R), and β-globin mRNA. Data are fit to exponential functions (lines on plots) used to derive mean times and rates (Table S1). **e**. Quantification of initiation complexes exhibiting different numbers of eIF1 reassociation events.

The 66 G_4_C_2_ repeats inhibited initiation after the scanning step. With both the native CUG site and the artificial AUG site, the duration of the initial eIF1 binding event (i.e., scanning) was 1.7 ± 0.1 s and 1.5 ± 0.1 s, respectively (Fig 3c). These durations are consistent with β-globin (2.0 ± 0.2 s) and our previous results with a short unstructured model RNA with an AUG start site^35^. In striking contrast, the subsequent window of time in which eIF1 transiently reassociated with the initiation complexes was 10-fold longer for mRNAs with 66 G_4_C_2_ repeats (113 ± 8.0 s (CUG) and 132 ± 16.0 s (AUG) versus 13 ± 2.1 s on β-globin; Fig 3d). Reassociation of eIF1 was also more frequent (∼70% of the mRNA-ribosome initiation complexes had 3 or more reassociations compared to ∼25% on β-globin) but dissociated similarly (0.9 s ± 0.1 s on 66-repeat and 0.5 s ± 0.1 s on AUG-66R mRNAs versus 1.1 ± 0.4 s on β-globin; Fig 3e-f: Fig S3a). The brief duration of eIF1 rebinding is consistent with a conformational transition of the initiation complex from a scanning-competent to scanning-arrested state, which introduces steric clashes that destabilizes eIF1 binding^30,70^. The normal kinetics of the initial eIF1 departure, along with the extended window for eIF1 rebinding, suggests that pathogenic G_4_C_2_ repeats impair initiation after start codon recognition.

### RAN translation initiation has commitment issues

After scanning and initial recognition of the start site, eIF5 directly competes with eIF1 to bind initiation complexes^29,34,35^. Productive eIF5 binding terminates eIF1 reassociation stimulating GTP hydrolysis and release of eIF2-GDP, P_i_ and eIF5 from the initiation complex^37,39^ and subsequent eIF5B association and 60S ribosomal subunit joining^36,71^. Given that the window of eIF1 reassociation was extended during initiation with 66 G_4_C_2_ repeat mRNAs, we hypothesized that the repeats inhibit eIF5 binding. We therefore applied our tRNA_i_-to-eIF5 FRET signal to monitor eIF5 directly during initiation. For these experiments, we assembled 43S initiation complexes with eIF2-Cy3-tRNA_i_-GTP ternary complex (green) with eIF5-Cy5.5 (purple) and unlabeled eIF1 and monitored Cy3-tRNA_i_ (the 43S complex), Cy3.5-eIF5B, and tRNA_i_-to-eIF5 FRET (purple; (Fig 4a-b)).

**Figure 4.**
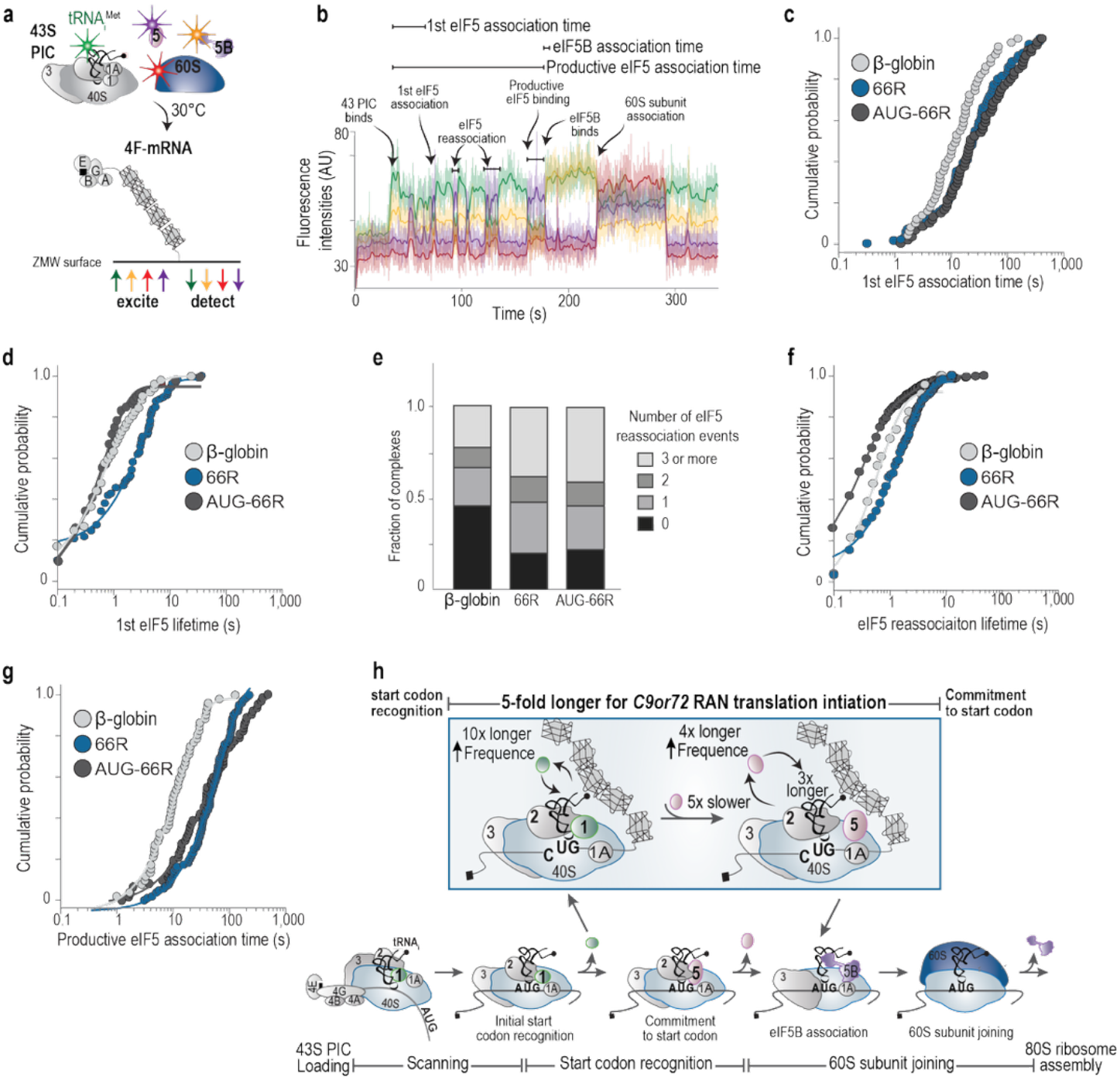
eIF5 dynamics are disrupted during C9 RAN translation initiation. **a**. Schematic of the post-start codon recognition assay. **b**, Representative traces tracking Cy3-tRNA_i_ (green), eIF5-Cy5.5 (purple), eIF5B-Cy3.5 (yellow), and 60S-Cy5 (red). eIF5 binding is detected via Cy3-Cy5.5 FRET (purple). Molecular binding events are annotated in the example trace. **c, e–g**. Cumulative probability plots of eIF5 dynamics for the 66 G4C2 repeat 66R mRNA, the 66R mRNA with a CUG to AUG mutation, and β-globin mRNA. **d**. Fraction of initiation complexes with the indicated number of eIF5 reassociation events. **h**. Proposed model of C9 RAN translation initiation illustrating altered eIF1 and eIF5 dynamics as revealed by single-molecule measurements. Fold changes observed for *C9orf72* RAN translation initiation relative to β-globin initiation are noted. Arrows indicate whether the fold change is an increase or decrease in the measured parameter.

The 66 G_4_C_2_ repeats and CUG start site perturbed eIF5 binding in multiple ways. In comparison to β-globin, the delay from loading of the 43S initiation complex onto the *C9orf72* RAN mRNA until eIF5 first bound was 5-fold longer (53 ± 13 s versus 10 ± 0.6 s; Fig 4c). While much slower to occur, this timing places the initial eIF5 binding event within the eIF1 rebinding window measured above, as expected. However, the duration of the initial eIF5 binding event was nearly 4-fold longer than on β-globin (5.4 ± 2.2 s versus 1.3 ± 0.5 s; Fig 4d). Subsequent eIF5 rebinding events were more frequent (∼40% of molecules had 3 or more reassociation events compared to 20% for β-globin) and had longer lifetimes (3.2 ± 0.3 s versus 0.7 ± 0.2 s; Fig 4e, 4f) than on β-globin. Despite the striking delay caused by the 66 G_4_C_2_ repeats, eIF5 ultimately bound (69 ± 3.4 s versus 14 ± 1.0 s on β-globin; Fig 4g) and progressed initiation complexes to eIF5B binding after eIF5 departure (2.1 ± 0.2 s on 66-repeat mRNA and 2.5 ± 0.3 s on β-globin; Fig S4). The duration of the final eIF5 binding event matched the duration observed on β-globin here and in our prior report (Fig S4b; ref). Substitution of the native CUG with a canonical AUG start site decreased the duration of the first eIF5 binding event by 5-fold to 1.2 s ± 0.2 s and accelerated eIF5B binding by 2-fold (1.0 ± 0.4 s Fig 4d, Fig S4c), consistent with increased activity with AUG start site in our bulk translation assays. This substitution did not alleviate the delays in the first and final productive eIF5 binding, suggesting this effect is a consequence of the 66 G_4_C_2_ repeats and not the identity of the start codon (Fig 4c, 4f).

We conclude that disease-length G_4_C_2_ repeats and the native CUG near-cognate start site act synergistically to induce a ribosomal state that inhibits eIF5 binding and activity, likely by funneling initiation complexes into unproductive, off-pathway states. Nevertheless, a subpopulation of initiation complexes ultimately resolves the inhibitory state and successfully initiates translation.

## Discussion

We define a mechanistic model of C9orf72 RAN translation initiation (Fig. 4h). It is well-established that both the G_4_C_2_ repeats and the upstream near-cognate CUG codon are necessary for *C9orf72* RAN translation^52,61^, but the mechanistic basis for their regulatory effects remained unclear. By monitoring individual ribosomal subunits and key eIFs as they progress through initiation in real time, we demonstrate that the G_4_C_2_ repeats promote *C9orf72* RAN translation by acting as a steric or thermodynamic barrier to initiation, agreeing with prior models ^54,55^. The G_4_C_2_ repeats extend the cumulative residency time of the initiation complex at the CUG start codon, which modulates eIF1 and eIF5 dynamics and underlies usage of this non-canonical start site.

We found that the 43S initiation complex is recruited and scans without impediment to the CUG codon on RNAs with 66 G_4_C_2_ repeats, using initial eIF1 departure as a proxy. The kinetics of scanning and initial start codon recognition in *C9orf72* RAN translation initiation match initiation on canonical mRNAs (this study and^35^). Without G_4_C_2_ repeats of sufficient length and structure positioned downstream, the CUG start site is bypassed by the scanning ribosomal complex. This suggests that the G_4_C_2_ repeats facilitate CUG initiation by arresting scanning at the CUG start site. The dramatic delay in subsequent eIF5B binding during *C9orf72* RAN translation initiation on RNAs with pathogenic-length G_4_C_2_ repeats indicates that the repeats also modulate steps after start codon recognition stages of initiation, ultimately increasing the cumulative time that the initiation complex spends at the CUG start site.

After reaching the CUG codon, initiation complexes assembled on G_4_C_2_-containing RNAs are in a state suboptimal for eIF5 binding and function. This state is characterized by multiple, nonproductive binding events by eIF5, suggesting that the conformational landscape of the 48S complex is distorted in a manner that disfavors eIF5 function. Eventually, this perturbed state resolves and a final productive eIF5 interaction with the initiation complex is followed by eIF5B recruitment and 60S subunit joining, both of which proceed with unperturbed kinetics. These results support a model in which the G_4_C_2_ repeats selectively destabilize the central GTP hydrolysis-dependent commitment step of initiation, acting after scanning and prior to eIF5B association and subunit joining. Resolution of the eIF5-impaired intermediate is rate-limiting for *C9orf72* RAN translation. This disruption is independent of start codon identity, suggesting that pathogenic-length G_4_C_2_ repeats alone are sufficient to modulate 48S complex dynamics at this step. However, the use of a near-cognate CUG start codon further impairs 60S subunit joining, indicating that start codon identity modulates later steps in initiation, possibly due to destabilized codon–anticodon pairing. Moreover, we observe synergistic effects between G_4_C_2_ repeat length, repeat proximity to the CUG, and codon identity, indicating that RNA structure and sequence context together trigger initiation and modulate initiation efficiency.

The nature of the RNA structure adopted by G_4_C_2_ repeats during translation remains an unresolved but critical issue. Both GQ and hairpin conformations have been implicated in RAN translation regulation^6,54,55^. In our assays, destabilizing GQ but not hairpin structures inhibited CUG initiation on 66R RNAs, supporting a functional role for GQ structures in promoting *C9orf72* RAN translation. Although hairpin structures with predicted free energy (ΔG) of ∼ –22 kcal/mol may also favor initiation at near-cognate sites^28^, the loss of translational activity upon GQ disruption points to a more specific requirement for GQ conformation. G_4_C_2_ repeat RNAs likely adopt metastable or dynamic structural ensembles during translation, potentially shifting between hairpin, GQ, or a heterogeneous mix of conformations^72-74^.

Our findings underscore the importance of RNA structure in modulating translation. How RNA remodeling factors, such as helicases, in *C9orf72* RAN translation is unclear^8,63,75^, and cellular stress^52,76-79^ modulate *C9orf72* RAN translation mechanisms remain key open questions. Future work should explore how these factors influence *C9orf72* RAN translation initiation and the structural landscape of repeat-containing RNAs. Our work and the accompanying manuscript by Kaufhald et al. redefine the model of *C9orf72* RAN translation initiation. We show that RNA G_4_C_2_ repeat structure and sequence context remodel initiation complexes after ribosomal scanning ends, generating a kinetic and conformational bottleneck in eIF5-mediated commitment. This model provides a mechanistic framework for understanding how repeat expansions modulate canonical translation pathways and may be broadly relevant to other repeat expansion disorders that involve RAN translation.

## Acknowledgements

We are grateful to members of the Lapointe, Gitler, and Puglisi labs for helpful discussions and feedback. We thank Peter Sarnow (Stanford) for sharing cell culture equipment, and Christopher S. Fraser (UC Davis) for initial guidance on factor purification.

## Author contributions

R.G. and J.D.P. conceived and designed study; R.G. acquired, analyzed data; R.G., J.D.P, C.P.L. interpreted data; S.B.Y., A.P., C.A. and M.Z.P. reagent preparation; C.P.L signal development and analysis code; R.G. drafted manuscript; J.D.P., A.D.G., C.P.L. manuscript writing and revisions; C.A. and M.Z.P, S.B.Y manuscript revision.

## Data, materials, and code availability

Raw single-molecule data were processed with custom MATLAB code. Code is available through GitHub: https://github.com/LapointeLab/eIF1-eIF5-2024-pape and https://github.com/LapointeLab/eIF5B-2022-paper. Source data will be made available on GitHub.

## Materials and Methods

*Plasmid construction and cloning*. Mammalian expression plasmids containing 2 and 66 G_4_C_2_ repeats flanked by native intronic sequence have been described^19^. These plasmids were used to construct plasmids appropriate for T7 in vitro transcription using restriction digestion and ligation cloning. The 5′ end of the leading intronic sequence has an upstream adjacent HindIII restriction site, and the 3′ intronic sequence contains a downstream adjacent NotI site adjacent to the G_4_C_2_ repeats. These sites were used to transfer the entire block to a vector between a 5′ T7 promoter sequence and a 3′ nLuc coding sequence. The nLuc coding sequence lacks an AUG start codon and contains a 3′ 30-nucleotide polyA tail after the stop codon followed by a SpeI restriction site for plasmid linearization. The leader sequence contains a native BssHII two nucleotides upstream of the first G of the repeats. To construct plasmids with modifications in the leader sequence, the HindIII and BssHII restriction sites were used to replace the leader sequence with a gene block (IDT, Newark, NJ, USA) containing the desired modification and the same flanking restriction sites. The same approach was used to construct the 6-repeat plasmid using the BssHII and NotI restriction site. For plasmids with the CUG to AUG and CUG to CCC mutations, and moving the nLuc CDS into alternative reading frames, site-directed mutagenesis (QuickChange Lightning, Agilent) was carried out on the 2 G_4_C_2_ repeat plasmid. In the resulting plasmids, the 2 G_4_C_2_ repeats were replaced with 66 G_4_C_2_ repeats using the restriction digestion-ligation approach. The CCC mutation also has a mutated Kozak sequence (Table S2). A separate 2 G_4_C_2_ repeat plasmid was constructed for use as the transcription template for the 2-repeat mRNA. It contains an AUG and Kozak sequence upstream of the nLuc CDS that was needed for detectable nLuc activity in in vitro translation (IVT) assays. Leader sequences are provided in Table S2. All enzymes used in cloning (restriction digest, ligation) were purchased from NEB and used according to the provided standard protocols. After confirming plasmid sequences (Sequetech, Mountain View, CA, USA or GeneWiz, South Plainfied, NJ, USA), plasmids were amplified and purified using Qiagen Giga Prep kits. Plasmid sequencing was reconfirmed after amplification, given the repeat instability even in Stbl3 cells (Thermo Scientific).

### mRNA preparation

The GAPDH and β-globin plasmids and mRNA preparation have been previously described^35,46,80^. The G_4_C_2_ repeat mRNAs used for nLuc activity assays were transcribed using Message MAX T7 ARCA-Capped Message kit (Cell Script) following a standard scaled-up protocol. Transcription reactions were supplemented with 2 M (final concentration) betaine to eliminate abortive transcription products and produce a single full-length mRNA. The mRNAs used in the 7-deaza-G IVT assays were transcribed with T7-FlashScribe kit (Cell Script). The mRNA transcription reactions contained 2 M betaine and either GTP or 7-deaza-GTP. These mRNAs were post-transcriptionally capped with Cap-1 (Vaccinia Capping kit, NEB). The mRNAs used for single-molecule experiments were similarly transcribed and post-transcriptionally capped, and then a biotin to was added to the 3^′^ end of the mRNAs by formation of a covalent hydrazone bond. The RNA terminal 3^′^ cis-diol was oxidized with sodium periodate to form an aldehyde and then reacted with Biotin-hydrzide to form the covalent hydrazone linkage following previously describe methods^81^.

Transcribed mRNAs were purified using MEGAclear (ThermoFisher). Folding of the RNA was done after the final purification immediately after elution from the spin column. The mRNA (concentration at or below 100 nM) was cooled to room temperature and then supplemented with KOAc to a final concentration of 100 mM. For IVT assays, the mRNAs were concentrated to 2.6 μM. mRNAs for single-molecule experiments were not concentrated. All repat-mRNAs containing more than 6 G_4_C_2_ repeats were stored at 4°C for no more than 10 days.

### nLuc activity assays

HeLa cytoplasmic extract in vitro translation assays. HeLa cell-free translation (ThermoFisher, #88884) reactions (10 μL) were programmed with 100 nM mRNA according to the kit protocol. Reactions were incubated at 37°C for 45 min and then assayed for nLuc luminescence (Promega nanoGlow kit and protocol) in a 384-well plate. Luminescence was measured at 3 min intervals for 30 min (BioTek Synergy Neo2 plate reader). The rate of signal decay was used to evaluate sample quality. Biological replicates (n) for experiments are noted in figure legends. Data analysis and visualization was done with GraphPad Prisim10 software.

In-gel detection of nLuc was done using the Promega in-gel detection kit following the standard protocol. Luminescence was imaged with a ChemiDoc MP Imager (BioRad).

Real-time nLuc activity assays were done with 25 μL reactions programed with 80 nM mRNA. A 1:10 v/v of nGlow substrate was added to the cell-free translation reactions. Before the addition of mRNA, the IVT reactions were transferred to non-adjacent wells in a 384-well plate and equilibrated for 10 min at 30°C in the plate reader. All other reagents were maintained at 30°C before being added to the IVT reactions. The order-of-addition is outlined in Fig S1g. Luminescence was monitored in situ (30°C). for 36 min, with 9 or 15 s intervals. Data was analyzed in MATLAB^82^.

### Biomolecules for single-molecule assays

Labeled ribosomal subunits. Human 40S and 60S ribosomal subunits were purified as described^83^. Ribosomal subunits from edited HEK293T cell lines containing either RPS15-ybbR or RPL5-ybbR were labeled with fluorescent dyes as previously described^46,80^.

Initiator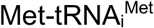. Met-tRNA_i_ was in vitro transcribed and aminoacylated as previously described^23,46,80^. Synthetic human Met-tRNA_i_ with U46 replaced with U-C6 Amino linker conjugated to Cy3 SE was purchased from TriLink (San Dieago, CA, USA). Aminoacylation has been detailed previously^35^. The amino acid charging efficiencies of Met-tRNA_i_ are > 90% based on acid urea PAGE analyses^35^.

Unlabeled eIFs. Recombinant proteins were purified as described in references: eIF4AI^84,85^, eIF4B^46^, eIF4G165-159^46^, eIF4E^85,86^, eIF1^84^, eIF1A^84^, eIF3j^84^, eIF5^23^, and eIF5B^46^,. Endogenous human eIF2 and eIF3 were purified as described^84,87^.

Labeled eIFs. Fluorescently labeled eIF1 C69A^35^, eIF5-ybbR^35^, and eIF5B-ybbR^46^ were purified and labeled as previously described.

### Real-time single-molecule assays preparation

All real-time imaging was carried out on a modified Pacific Biosciences RSII DNA sequencer with Maggie software (v. 2.3.0.3.154799)^88^. Single illumination (532 nm laser at 0.32 µW/µm2) excitation of Cy3 and Cy3.5, or dual illumination (532 nm laser and 642 nm laser, 0.1 µW/µm2) excitation of Cy5 and Cy5.5 fluorophores. Four-color fluorescence emission was detected at 10 frames per second for 800 s. Zero-mode waveguides (ZMW) chips (Pacific Biosciences) with PEG-biotin conjugated to the surface were washed with 0.2% Tween-20 and TP50 buffer (50 mM Tris-OAc pH 7.5, 100 mM KCl), and then incubated for 5 min with 1 μM neutravidin and surface passivating agents (0.7 mg/mL Ultrapure BSA, and 1.3 µM of pre-annealed DNA blocking oligos^89^ in TP50 buffer. The imaging surface was washed three times with TP50 buffer, and then 2 times with initiation reconstitution (REC) buffer (20 mM HEPES-KOH, pH 7.5, 70 mM KOAc, 2.5 mM Mg(OAc)_2_, 0.25 mM spermidine, 0.2 mg/mL creatine phosphokinase, 1 mM ATP•Mg(OAc)_2_, and 1 mM GTP•Mg(OAc)_2_. All single-molecule fluorescence experiments were performed as described previously^35,46,80^.

eIF2–GTP– 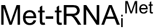 ternary complex (TC): 3.3 µM eIF2 was incubated in REC buffer (no ATP•Mg(OAc)_2_) for 10 min at 37 °C and then incubated with 2.3 µM of unlabeled 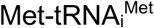 or 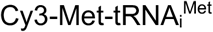for 5 min at 37°C.

43S pre-initiation complex (43S PIC): 1 µM eIF1, 1 µM eIF1A, 500 nM TC (by eIF2), 1 µM eIF5, 400 nM eIF3, 1.2 µM eIF3j, and 240 nM 40S or Cy3-40S subunits (≊10 nM pre-initiation complex) were incubated for 5 min at 37°C in REC buffer. Fluorescently labeled factors were used at the same concentration as unlabeled factors during pre-initiation complex formation.

### Real-time single-molecule initiation assays

5^′^ m^7^G capped and 3^′^-biotinylated mRNA was tethered to the imaging surface (10 min incubation at room temperature). After tethering, the chip surface was washed 2 times with REC buffer, and then 2 times with imaging buffer^90,91^ (REC buffer plus 62.5 µg/mL casein, 5 mM TSY, 2 mM protocatechuic acid, and 0.06 U/µL protocatechuate-3,4-dioxygenase). A solution containing of 2 µM eIF4A, 440 nM eIF4B, 260 nM eIF4G, and 320 nM eIF4E in 20 µL of imaging buffer was added to the mRNA tethered to the ZMW surface and incubated at room temperature for ∼10 minutes. At the start of data acquisition, a solution containing the 43S PIC, eIF1, eIF5, Cy3.5-eIF5B, and 60S subunits (unlabeled or labeled with Cy5.5) was delivered to the surface of the ZMW chip. The final concentrations after delivery were as follows: 1 µM eIF4A, 220 nM eIF4B, 230 nM eIF4G, 320 nM eIF4E, 10 nM pre-initiation complex (by 40S subunits), 290 nM eIF1A, 290 nM eIF1 (unlabeled), and 290 nM eIF5 (unlabeled). Cy5-eIF1 and Cy5.5-eIF5 replaced the unlabeled proteins in the 43S pre-initiation complex formation when required, and no additional protein was added to the delivery solution. The final concentration of Cy5-eIF1 and Cy5.5-eIF5 was 40 μM in these experiments. In some experiments using Cy3-40S subunits and Cy3.5-5B, Cy5-60S subunits were used in place of Cy5.5-60S subunits (noted in Sup Table1).

### Data analyses

Fluorescence intensities were recorded for the duration of the experiment. The resulting movies were processed and converted to single-molecule traces using MATLAB R2019a as described previously^88^. Manual state (fluorescence on or off) assignment of each fluorescent signal and subsequent statistical analysis was done using custom MATLAB scripts.

To determine association and dissociation kinetics, the timing of the molecular binding events from single molecules (fluorescence traces from a single mRNA immobilized in a ZMW well) was used to calculate cumulative probability functions of the observed data (MATLAB cdfcalc)^92^. The cumulative probability functions were fit to single-or double-exponential functions (MATLAB cftool):

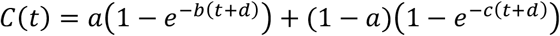

Where a is the amplitude or fraction of the molecules, t is time (s), b and c are rates, and d is an adjustment factor. A single-exponential fit was used when a ≥ 0.9.

Fluorescence traces in the main text figures were manually corrected to account for spectral bleed-through of the fluorescence signals. Reported errors for derived rates and constants represent 95% C.I. yielded from fits to linear, single-exponential, or double-exponential functions, as indicated.

