## Supplemental Figures for "RNA regulates repeat-associated non-AUG (RAN) translation initiation in *C9orf72* FTD/ALS"

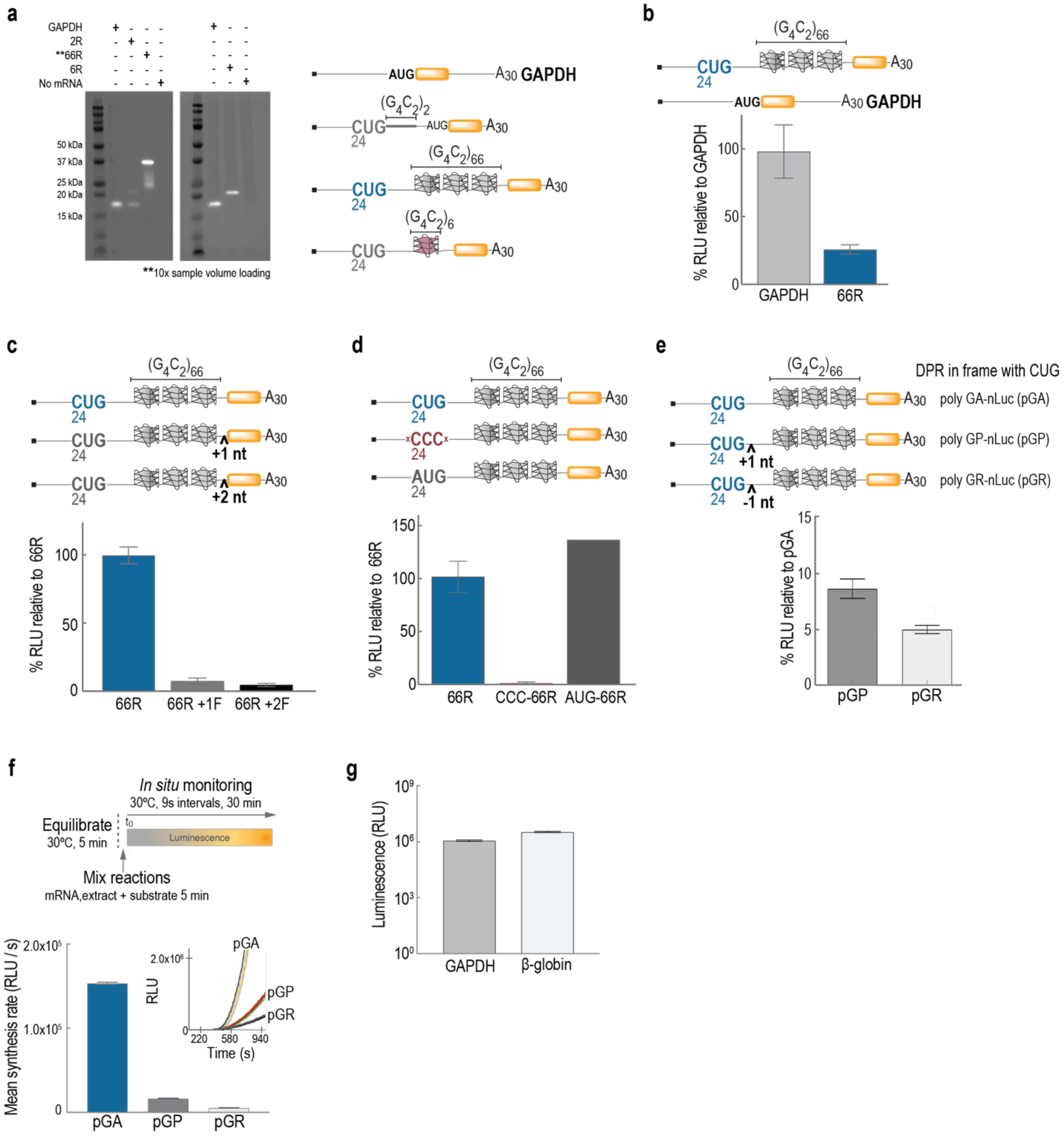


**Supplemental Figure 1: *In vitro* translation (IVT) of mRNA reporters in HeLa cytoplasmic extract recapitulates *C9orf72* RAN translation.**

**a.** Translation of dipeptide repeat proteins (DPRs) is repeat length dependent. Samples from IVT reactions programmed with the mRNAs schematized on the right of the gel, were analyzed by SDS-PAGE. The number of $G_4C_2$ repeats (R) is noted for each reporter. In-gel detection of nLuc luminescence provides visualization of the synthesized protein size. The GAPDH coding sequence (CDS) was replaced with the nLuc coding sequence with an AUG start codon. The 2R reporter also has an nLuc AUG start codon. All other reporters lack an AUG start codon.

**b-e.** Relative nLuc activity of mRNA reporters schematized after IVT in the panel. Error bars represent standard deviation (n = 3 biological replicates except where noted). See Supplemental Table 2 for nt substitutions the native leader sequence upstream of the repeats where applicable.

**b.** Level of *C9orf72* RAN translation relative to a control cellular mRNA.
**c.** *C9orf72* RAN translation in multiple reading frames relative to in-frame translation. Single or double nucleotide (nt) insertions upstream of the nLuc CDS were used to change the nLuc reading frame (RF).
**d.** *C9orf72* RAN translation in-frame translation initiates at a CUG codon 24 nts upstream of first G of the repeats. The CUG codon was substituted with and AUG or CCC codon at this position. The CCC construct also has a disrupted Kozak context sequence (n = 4).
**e**. DPR identity modulates *C9orf72* RAN translation level. Translation level of poly-glycine-proline (pGP) and poly-glycine-arginine (pGR) relative to poly-glycine-alanine (pGA) is plotted. A single nt insertion or deletion upstream of the G4C2 repeats was used to change the identity of the in-frame DPR.
**f.** DPR identity modulates *C9orf72* RAN translation rate. *In situ* monitoring of nLuc activity in real time was used to determine the synthesis. Inset in bar plot is the nLuc luminescence accumulation in real time.
**g.** Translation level of control mRNAs used in bulk and single-molecule experiments.

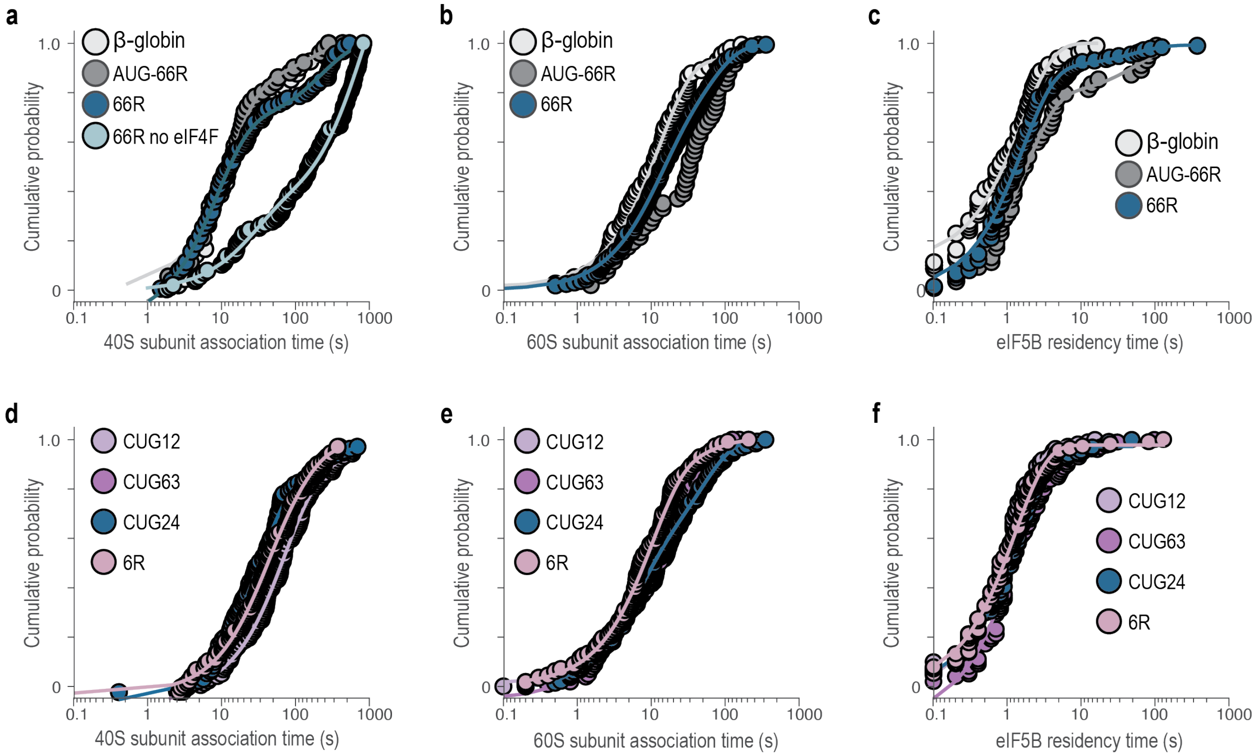


**Supplemental Figure 2: Initiation dynamics during *C9orf72* RAN translation initiation.** Cumulative probability plots showing binding dynamics of the 40S-Cy3 ribosome, eIF5B-Cy3.5, and 60S-Cy5. See Figure 2a-b for experimental schematic and definition of molecular biding parameter in the plots.

**a-c.** Molecular binding dynamics on 66 $G_4C_2$ repeat (66R), 66R with a CUG to AUG mutation (AUG-66R), and β-globin mRNAs.

**d-f.** Molecular binding dynamics on 66R mRNAs with 12-nt (CUG12), 24-nt (CUG24), or 63-nt (CUG63) spacing between the CUG start codon and the $G_4C_2$ repeats, and the 6R mRNA.

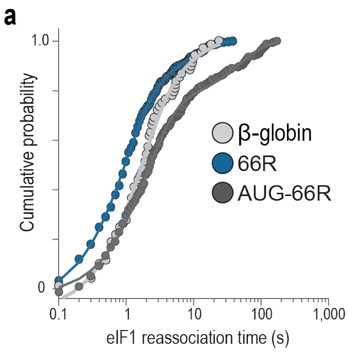


**Supplemental Figure 3: Reassociation time of eIF1 after start codon recognition during *C9orf72* RAN translation initiation.** Cumulative probability of eIF1 reassociation time on 66 $G_4C_2$ repeat (66R), 66R with a CUG to AUG mutation (AUG-66R), and β-globin mRNAs. See Figure 3a-b for experimental schematic and definition of eIF1 reassociation.

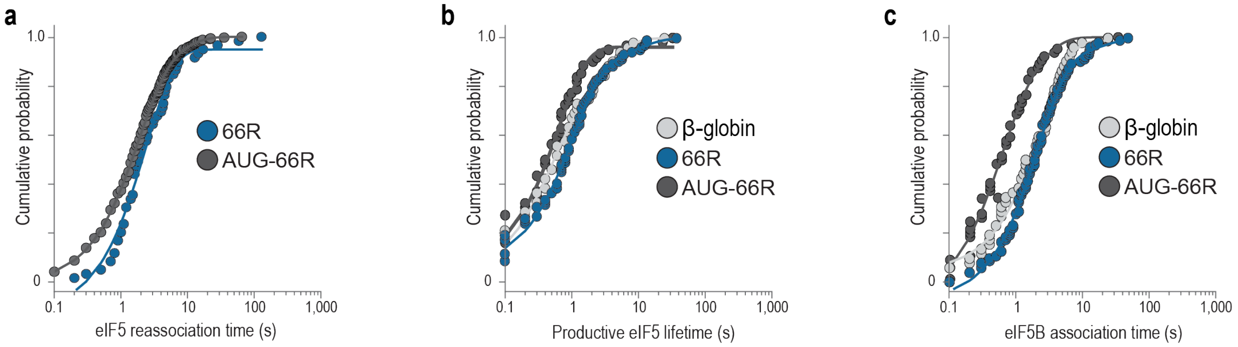


**Supplemental Figure 4: eIF5 and eIF5B dynamics during *C9orf72* RAN translation initiation.** Cumulative probability of molecular binding events on 66 $G_4C_2$ repeat (66R), 66R with a CUG to AUG mutation (AUG-66R), and β-globin mRNAs. See Figure 4a-b for experimental schematic and definition of molecular events.

**a.** Reassociation time of eIF5.

**b.** Lifetime of the final eIF5 binding event that precedes eIF5B association.

**c.** Association time of eIF5B following the departure of the final eF5 binding event.
